# Constitutive activation of leucine-rich repeat receptor kinase signaling pathways by BAK1-interacting receptor-like kinase 3 chimera

**DOI:** 10.1101/2020.02.18.954479

**Authors:** Ulrich Hohmann, Priya Ramakrishna, Kai Wang, Laura Lorenzo-Orts, Joel Nicolet, Agnes Henschen, Marie Barberon, Martin Bayer, Michael Hothorn

**Author notes:** Institute of Molecular Biotechnology of the Austrian Academy of Sciences (IMBA) & Research Institute of Molecular Pathology (IMP), Vienna Biocenter (VBC), 1030 Vienna, Austria. Research Institute of Molecular Pathology (IMP), Vienna Biocenter (VBC), 1030 Vienna, Austria. The author(s) responsible for distribution of materials integral to the findings presented in this article in accordance with the policy described in the Instructions for Authors (www.plantcell.org) are: Martin Bayer and Michael Hothorn.

## Abstract

Receptor kinases with extracellular leucine-rich repeat domains (LRR-RKs) form the largest group of membrane signaling proteins in plants. LRR-RKs can sense small molecule, peptide or protein ligands, and may be activated by ligand-induced interaction with a shape complementary SOMATIC EMBRYOGENESIS RECEPTOR-LIKE KINASE (SERK) co-receptor kinase. We have previously shown that SERKs can also form constitutive, ligand-independent complexes with the LRR ectodomains of BAK1-interacting receptor-like kinase 3 (BIR3) receptor pseudokinases, negative regulators of LRR-RK signaling. Here we report that receptor chimaera in which the extracellular LRR domain of BIR3 is fused to the cytoplasmic kinase domains of the SERK-dependent LRR-RKs BRASSINOSTEROID INSENSITIVE1, HAESA and ERECTA form tight complexes with endogenous SERK co-receptors in the absence of ligand stimulus. Expression of these chimaera under the control of the endogenous promoter of the respective LRR-RK leads to strong gain-of-function brassinosteroid, floral abscission and stomatal patterning phenotypes, respectively. Importantly, a BIR3-GSO1/SGN3 chimera can partially complement *sgn3* Casparian strip formation phenotypes, suggesting that GSO1/SGN3 receptor activation is also mediated by SERK proteins. Collectively, our protein engineering approach may be used to elucidate the physiological functions of orphan LRR-RKs and to identify their receptor activation mechanism in single transgenic lines.

## Introduction

Plant-unique membrane receptor kinases characterized by an extracellular domain, a single membrane spanning helix and a cytoplasmic dual-specificity kinase domain control many aspects of plant growth and development, form the first layer of the plant immune system and mediate symbiotic interactions (Hohmann et al., 2017). LRR-RKs form the largest class of receptor kinases known in plants (Shiu and Bleecker, 2001). Members of the family have been shown to sense small molecule (Wang et al., 2001), peptide (Gómez-Gómez and Boller, 2000; Matsubayashi, 2014; Santiago et al., 2016) and protein ligands (Huang et al., 2016; Lin et al., 2017; Zhang et al., 2017).

Brassinosteroids, whose biosynthesis involves the steroid 5ɑ steroid reductase DE-ETIOLATED2 (DET2) (Chory et al., 1991; Noguchi et al., 1999), are sensed by the ectodomain of the LRR-RK BRASSINOSTEROID INSENSITIVE1 (BRI1) with nanomolar affinity (Wang et al., 2001; Hothorn et al., 2011; Hohmann et al., 2018b). Brassinosteroid binding to the BRI1 ectodomain triggers the interaction with the LRR domain of a SOMATIC EMBRYOGENESIS RECEPTOR LIKE KINASE (SERK) co-receptor (Hothorn et al., 2011; She et al., 2011; Santiago et al., 2013; Sun et al., 2013; Hohmann et al., 2018b). Formation of this heterodimeric complex at the cell surface promotes interaction and trans-phosphorylation of the receptor and co-receptor kinase domains inside the cell (Wang et al., 2008; Bojar et al., 2014; Hohmann et al., 2018b; Perraki et al., 2018). BRI1 receptor activation triggers a cytoplasmic signaling cascade, which ultimately results in the dephosphorylation and activation of a family of basic helix-loop-helix transcription factors, including BRASSINAZOLE-RESISTANT1 (BZR1) and BRI1-EMS-SUPPRESSOR1 (BES1) (Wang et al., 2002; Yin et al., 2002; Vert and Chory, 2006; Nosaki et al., 2018). In *bes1-D* plants BES1 proline 233 is found replaced by leucine, which leads to constitutive brassinosteroid signaling responses by enhancing protein phosphatase 2A mediated dephosphosrylation (Yin et al., 2002; Tang et al., 2011).

The plant-unique SERK co-receptor dependent activation mechanism is conserved among many LRR-RKs (Hohmann et al., 2017), including the LRR-RK HAESA, which, for example, controls floral organ abscission in Arabidopsis by interacting with the peptide hormone IDA (Jinn et al., 2000; Meng et al., 2016; Santiago et al., 2016; Hohmann et al., 2018b). A SERK-dependent MAP kinase signaling pathway (Meng et al., 2015) with diverse roles in plant development involves the LRR-RK ERECTA (ER) and its paralogs ERECTA-LIKE1 (ERL1) and ERL2 (Torii et al., 1996; Shpak, 2013). ER, ERL1 and ERL2 together control stomata development and their correct spacing on the leaf surface (Shpak et al., 2005). Cysteine-rich EPIDERMAL PATTERNING FACTOR peptides (EPFs) bind to the ectodomains of ER, ERL1 and ERL2 which form constitutive complexes with the ectodomain of the receptor-like protein (RLP) TOO MANY MOUTH (TMM) (Yang and Sack, 1995; Nadeau and Sack, 2002; Lee et al., 2012, 2015; Lin et al., 2017). EPF peptide binding to these LRR-RK/LRR-RLP complexes triggers their interaction with SERK co-receptor kinases (Meng et al., 2015; Lin et al., 2017), which in turn leads to the initiation of a MAP kinase signaling pathway that includes the MAP3K YODA (Bergmann et al., 2004). Stimulation of the ERECTA pathway negatively regulates stomata formation (Lampard et al., 2009).

Complex structures and quantitative biochemical comparisons of different ligand-activated LRR-RK – SERK complexes have revealed a structurally and functionally conserved activation mechanism, relying on the interaction of the ligand bound receptor LRR ectodomain with the shape-complementary ectodomain of the SERK co-receptor (Santiago et al., 2013; Wang et al., 2015; Santiago et al., 2016; Hohmann et al., 2017; Lin et al., 2017; Hohmann et al., 2018b). The ligand binding specificity of plant LRR-RKs is encoded in their LRR ectodomains (Hohmann et al., 2017; Okuda et al., 2020). The kinase domain of the receptor, not of the SERK co-receptor, confers cytoplasmic signaling specificity (Hohmann et al., 2018b; Chen et al., 2019; Zheng et al., 2019). Recent genetic, biochemical and structural evidence suggest that not all plant LRR-RKs rely on SERKs as essential co-receptor kinases (Hu et al., 2018; Cui et al., 2018; Anne et al., 2018; Smakowska-Luzan et al., 2018; Zhang et al., 2017).

Protein engineering approaches have been previously employed to dissect the LRR-RK receptor activation *in planta*: A fusion protein combining the extracellular and trans-membrane domains of BRI1 (outerBRI1, oBRI1) with the cytoplasmic kinase domain of the rice immune receptor XA21 (innerXA21, iXA21) could trigger immune signaling in rice cells upon stimulation with brassinisteroids (He et al., 2000). It is now known that both BRI1 and XA21 rely on SERK co-receptor kinases for receptor activation (Li et al., 2002; Nam and Li, 2002; Santiago et al., 2013; Hohmann et al., 2018b; Chen et al., 2014). The heteromeric nature of LRR-RK – SERK complexes has been validated *in planta* using similar protein engineering approaches. Co-expression of a protein chimera of the immune receptor FLAGELLIN SENSITIVE 2 (FLS2) and its co-receptor SERK3 (oFLS2-iSERK3) with an oSERK3-iFLS2 construct led to immune signaling after stimulation with the FLS2 ligand flg22 in a transient expression system (Albert et al., 2013). Stable transgenic lines co-expressing oBRI1-iSERK3 and oSERK3-iBRI1 construct could partially rescue the BRI1 loss-of-function mutant *bri1-301* (Hohmann et al., 2018b).

The signaling specificity of the cytoplasmic kinase domain of LRR-RKs has been dissected using an oBRI-iHAESA chimera, which rescued the floral abscission phenotypes when expressed under the control of the HAESA promoter in the *haesa hsl2* double mutant (Hohmann et al., 2018b). A similar approach has been recently used to demonstrate that the LRR-RKs BRI1 and EMS1 share a common cytoplasmic signaling cascade (Zheng et al., 2019). However, all these approaches rely on ligand stimulus.

Recently, a constitutive, ligand-independent interaction between the LRR ectodomains of SERKs and of BAK1-INTERACTING RECEPTOR-LIKE KINASEs (BIRs) has been reported (Ma et al., 2017; Hohmann et al., 2018a). While BIR1 appears to have a catalytically active cytoplasmic kinase domain, BIR2-4 are receptor pseudokinases (Gao et al., 2009; Wang et al., 2011; Blaum et al., 2014). Different BIRs have been characterized as negative regulators of plant immune, floral abscission and brassinosteroid signaling (Gao et al., 2009; Halter et al., 2014; Leslie et al., 2010; Imkampe et al., 2017). Structural and biochemical analyses now implicate BIR proteins as general negative regulators of SERK co-receptor mediated LRR-RK signaling pathways (Moussu and Santiago, 2019). The ectodomains of BIR1-4 bind to SERK ectodomains with dissociation constants in the low micromolar range and target a surface area in SERKs normally required for the interaction with ligand-bound LRR-RKs (Hohmann et al., 2018b; Ma et al., 2017; Hohmann et al., 2018a). Thus, BIRs can efficiently compete with LRR-RKs for SERK binding, negatively regulating LRR-RK signaling pathways. In line with this, the elongated (*elg*) allele in SERK3, which weakens the interaction with BIRs but not with BRI1 results in a brassinosteroid-specific gain-of-function signaling phenotype, as BRI1 can more efficiently compete with BIRs for co-receptor binding (Jaillais et al., 2011; Hohmann et al., 2018a). Structure-guided mutations in the BIR – SERK ectodomain complex interface (BIR3 residues Phe146-Ala/Arg170-Ala) efficiently disrupt BIR – SERK signaling complexes *in vitro* and *in planta* (Hohmann et al., 2018a). Here we present protein fusions of the BIR3 LRR ectodomain and transmembrane helix (oBIR3) with the cytoplasmic domains of different SERK-dependent LRR-RKs (iBRI1, iHAESA, iER, iFLS2). Expressing these chimera under the control of endogenous/context-specific promoters, we obtain strong gain-of-function phenotypes for different LRR-RK triggered developmental signaling pathways. In addition, an oBIR3-iGSO1/SGN3 chimera supports a SERK-dependent activation mechanism for the LRR-RK GASSHO1/SCHENGEN3 in Casparian strip formation (Pfister et al., 2014; Okuda et al., 2020). Our strategy allows for the identification of gain-of-function phenotypes of orphan LRR-RKs whose ligands are unknown, and enables the elucidation of their receptor activation mechanism.

## Results

We compared the structure of a previously reported BRI1 – brassinolide (BL) – SERK1 complex (Protein Data Bank ID 4SLX, http://rcsb.org) with the recently reported complex structure of a BIR3 – SERK1 complex (PDB-ID 6FG8) (Santiago et al., 2013; Hohmann et al., 2018a). The BRI1 and BIR3 ectodomains bind SERK1 using non-identical but overlapping binding surfaces (Figure 1A). As in the BRI1 – SERK1 complex, the C-termini of BIR3 and SERK1 are in close proximity in the complex structure (Figure 1B,C). Based on the structural similarities, we generated an oBIR3 – iBRI1 chimera, in which the BIR3 ectodomain and trans-membrane helix are connected to the cytoplasmic domain of BRI1 (Figure 1D) (see Methods).

**Figure 1.**
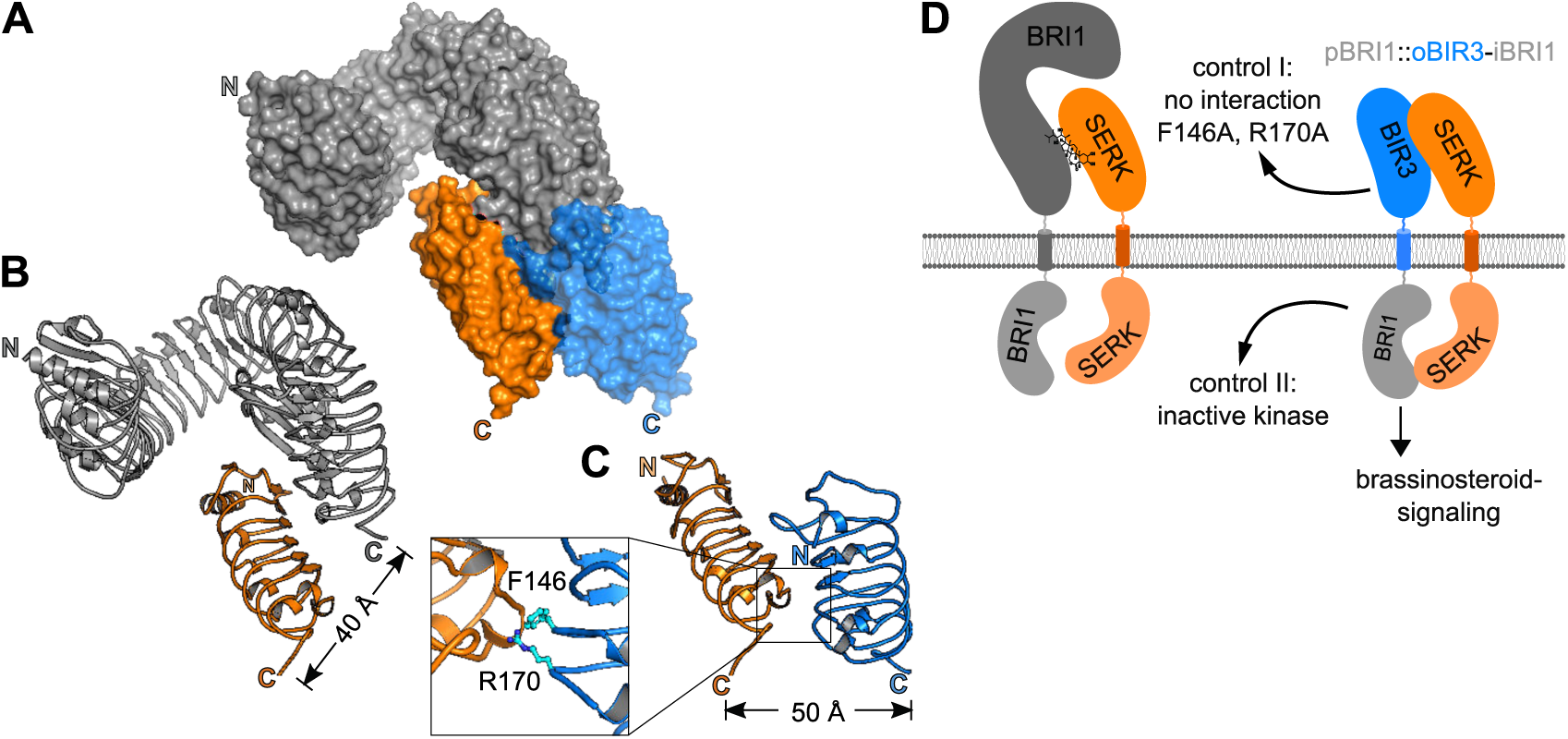
Structural overview BRI1 – SERK and BIR3 - SERK complexes. **(A)** Surface view of a structural superposition of a BRI1 – SERK1 (ectodomains shown in gray and orange, respectively; Protein Data Bank [PDB] – ID: 4LSX, http://www.rcsb.org/) and SERK1 – BIR3 (orange and blue; PDB-ID: 6FG8). The two structures are aligned on SERK1 (r.m.s.d. [root-mean-square deviation] = ∼ 0.3 Å comparing 143 corresponding C_ɑ_ atoms). **(B, C)** Ribbon diagrams of the BRI1 – SERK1 (B) and the BIR3 – SERK1 (C) complexes, with SERK1 shown in the same orientation. The distances between the respective C-termini are indicated (colors as in A). Inset: Close-up view of the BIR3 – SERK1 complex interface, with the interface residues Phe146 and Arg170 highlighted in bonds representation. Mutation of both residues to alanine disrupts the BIR3 – SERK1 complex *in vitro* and *in vivo* (Hohmann et al., 2018a). **(D)** Schematic overview of an entire BRI1 – BL – SERK signaling complex and the envisioned oBIR3-iBRI1 – SERK interaction.

We expressed the oBIR3 – iBRI1 fused to a C-terminal mCitrine (mCit) fluorescent protein tag under the control of the *pBRI1* promoter in a previously characterized *bri1* null mutant (Jaillais et al., 2011). oBIR3^F146A/R170A^ – iBRI1 and oBIR3 – iBRI1^D1027N^, which block BIR – SERK complex formation (Hohmann et al., 2018a) and BRI1 kinase activity (Bojar et al., 2014; Hohmann et al., 2018b), respectively, were used as controls. Independent oBIR3 – iBRI1 transgenic lines, but none of the control lines displayed the wavy hypocotyl phenotype characteristic of gain-of-function brassinosteroid mutants (Figure 2 A). Importantly, the wavy hypocotyl phenotype observed in oBIR3 – iBRI1 lines was also visible when plants were grown in the presence of the brassinosteroid biosynthesis inhibitor brassinazole (BZR) (Asami et al., 2000) (Figure 2A). This suggests that oBIR3 – iBRI1 triggered brassinosteroid signaling does not depend on endogenous brassinosteroids (Figure 2A). Consequently, we found all oBIR3-iBRI1 but none of the control lines to be constitutively active when expressed in the *det2* background (Chory et al., 1991), in which brassinosteroid levels are reduced (Fujioka et al., 1997) (Figure 2A). Quantification of three independent oBIR3-iBRI1 T3 lines revealed strong gain of function phenotypes, which are even more pronounced than the previously reported phenotype of the constitutively active *bes1-1D* mutant (Figure 2A) (Yin et al., 2002). Consistent with a constitutive activation of the brassinosteroid signaling, we found BES1 to be dephosphorylated in oBIR3 – iBRI1 but not in the control lines (Figure 2B), and BES1 dephosphorylation to also take place in the *det2* background (Figure 2C). We next performed co-immunoprecipitation (co-IP) experiments in our stable lines and found oBIR3 – iBRI1 and oBIR3 – iBRI1^D1027N^ to efficiently interact with the endogenous SERK3 co-receptor *in vivo*, while the oBIR3^F146A/R170A^ – iBRI1 control, which disrupts the interaction of the isolated BIR3 and SERK1/3 ectodomain *in vitro* (Hohmann et al., 2018a), could no longer bind SERK3 *in vivo* (Figure 2D). Taken together, the BIR3 ectodomain can promote brassinosteroid independent interaction with SERK3, and possibly other SERKs *in vivo*, resulting in a constitutive activation of the brassinosteroid signaling pathway. The control lines further suggest that this signaling complex is formed and stabilized by the ectodomains of BIR3 and SERK3, and requires the catalytic activity of the BRI1 kinase domain for signaling (Figure 2A).

**Figure 2.**
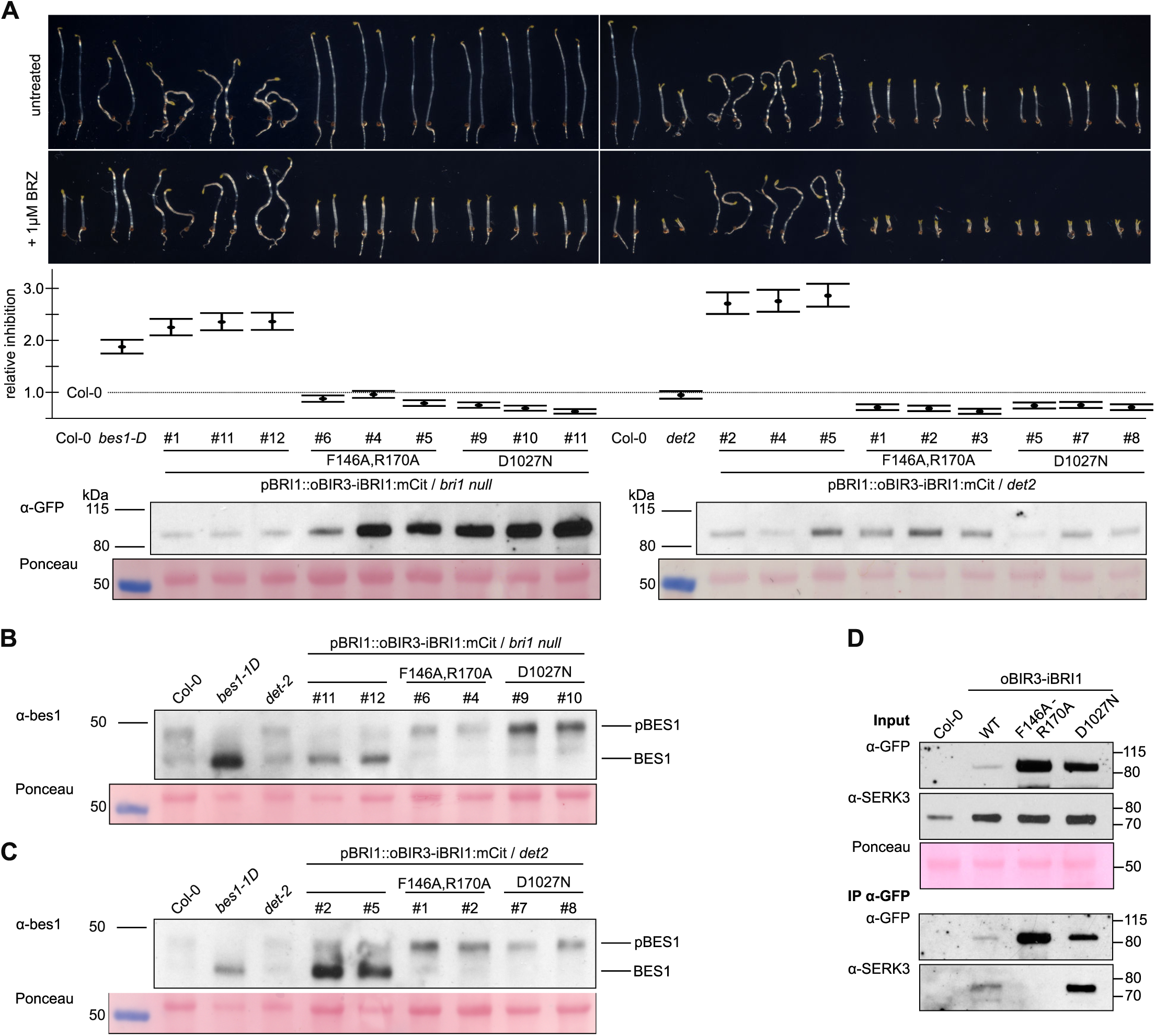
oBIR3 – iBRI1 chimera constitutively activate brassinosteroid signaling. **(A)** Hypocotyl growth assay of dark grown seedlings in the presence and absence of the BR biosynthesis inhibitor brassinazole (BRZ). Representative seedlings are shown in the top panel, with the quantification of the data (relative inhibition of hypocotyl growth in the presence of BRZ plotted together with lower and upper confidence intervals) below. For each sample n = 50 hypocotyls from 5 different ½ MS plates were measured. # numbers indicate independent lines. An anti-GFP westernblot together with the Ponceau – stained membrane as loading control, is shown alongside. **(B, C)** Anti-BES1 western blot on oBIR3-iBRI chimera in *bri1*-null (B) and *det2* (C) backgrounds, with the Ponceau – stained membranes shown alongside. **(D)** Co-immunoprecipitation experiment of oBIR3-iBRI1 chimera and SERK3. Shown alongside are the input western blots as well as a Ponceau–stained membrane.

We next tested if BIR3-based protein chimera can be used to activate a functionally distinct LRR-RK signaling pathway. The LRR-RK HAESA (HAE) shares the overall structure and activation mechanism with BRI1 (Santiago et al., 2013, 2016; Hohmann et al., 2018b), but the two receptors control very different developmental processes (Li and Chory, 1997; Jinn et al., 2000). We expressed an oBIR3-iHAE fusion protein (Figure 3A) fused to a C-terminal mCit tag under the control of the *pHAE* promoter in the *hae hsl2* mutant, in which floral organ abscission is delayed (Stenvik et al., 2008). We observed that expression of oBIR3-iHAE but none of the control lines rescued the floral abscission phenotype of the *hae hsl2* mutant (Figure 3B, C). Consistently, we found oBIR3 – iHAE and oBIR3 – iHAE^D1027N^ but not oBIR3^F146A/R170A^ – iHAE to interact with SERK3 in co-IP assays (Figure 3D).

**Figure 3.**
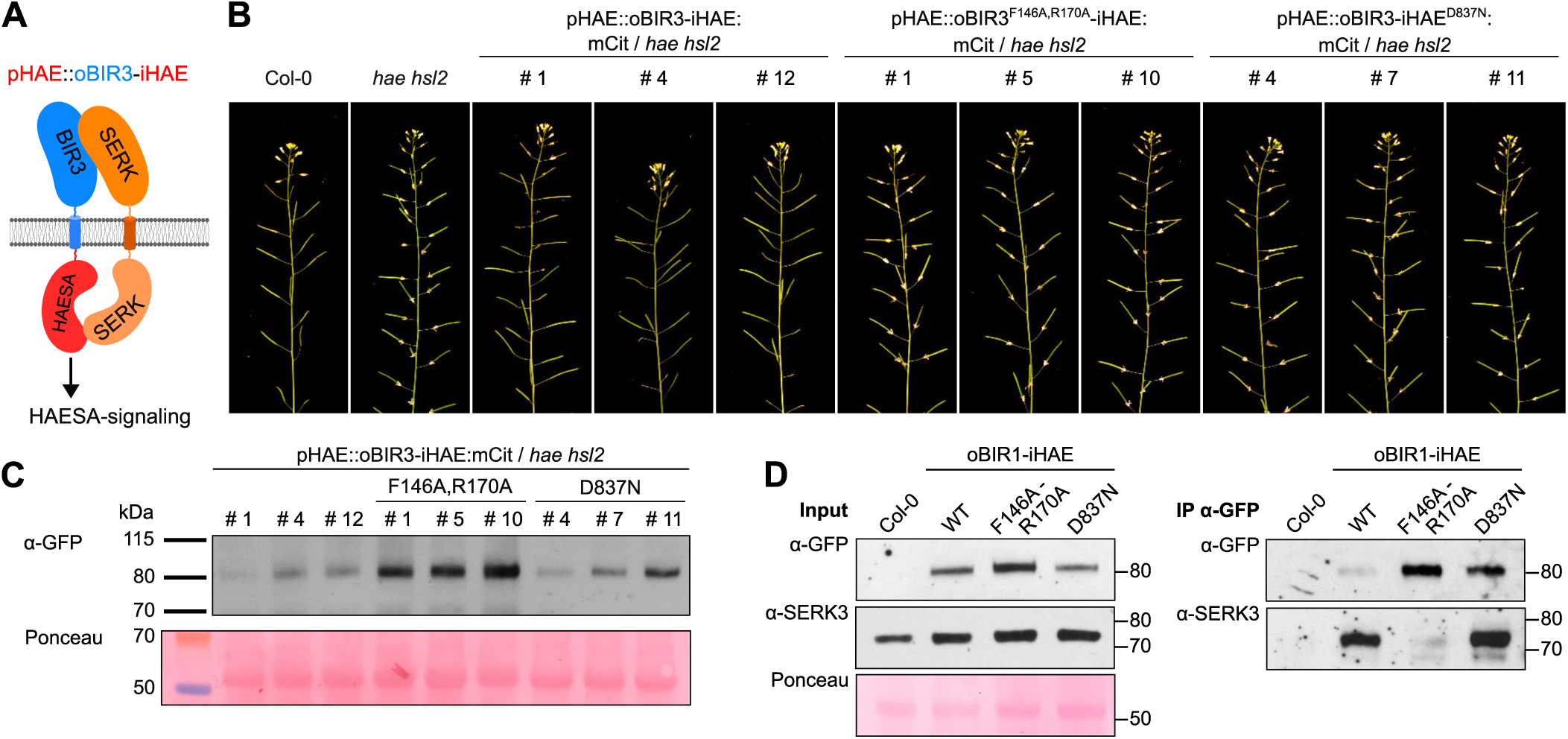
oBIR3 – iHAE chimera restore floral organ shedding in *hae hsl2* mutant plants. **(A)** Cartoon representation of the oBIR3-iHAE chimera. **(B)** Representative inflorescences of ∼9 week old Arabidopsis Col-0, *hae hsl2* and oBIR3-iHAE chimera; # numbers indicate independent lines. **(C)** Anti-GFP western blot together with the Ponceau–stained membrane as loading control. **(D)** Co-immunoprecipitation experiment of oBIR3-iHAE chimera and SERK3. Shown alongside are the input western blots as well as a Ponceau – stained membrane (left).

SERK proteins have been previously shown to allow for receptor activation of ERECTA family receptor kinases in protoderm formation and stomatal patterning (Meng et al., 2015). ER forms constitutive complexes with the LRR-RLP TMM to sense EPF peptides in stomatal patterning (Yang and Sack, 1995; Nadeau and Sack, 2002; Lee et al., 2012, 2015; Lin et al., 2017), but it is not understood at the mechanistic level how SERK co-receptor kinases allow for receptor activation of this LRR-RK/LRR-RLP signaling complex (Lin et al., 2017). To test if the receptor activation mechanism is conserved among BRI1, HAESA and ER, we expressed a chimeric oBIR3-iER construct fused to a C-terminal yellow fluorescent protein YPET specifically in the stomata lineage under control of the meristemoid-specific *MUTE* promoter (Figure 4A) (Pillitteri et al., 2007). Previous experiments demonstrated that constitutive activation of the ERECTA pathway in differentiating meristemoids leads to developmental arrest of guard mother cells (Lampard et al., 2009). To test the signaling specificity of our oBIR3-iER chimera, we also expressed a chimeric fusion of the innate immunity receptor FLS2 (Gómez-Gómez and Boller, 2000) under the control of the *MUTE* promoter (oBIR3-iFLS2-YPET) (Figure 4A).

**Figure 4.**
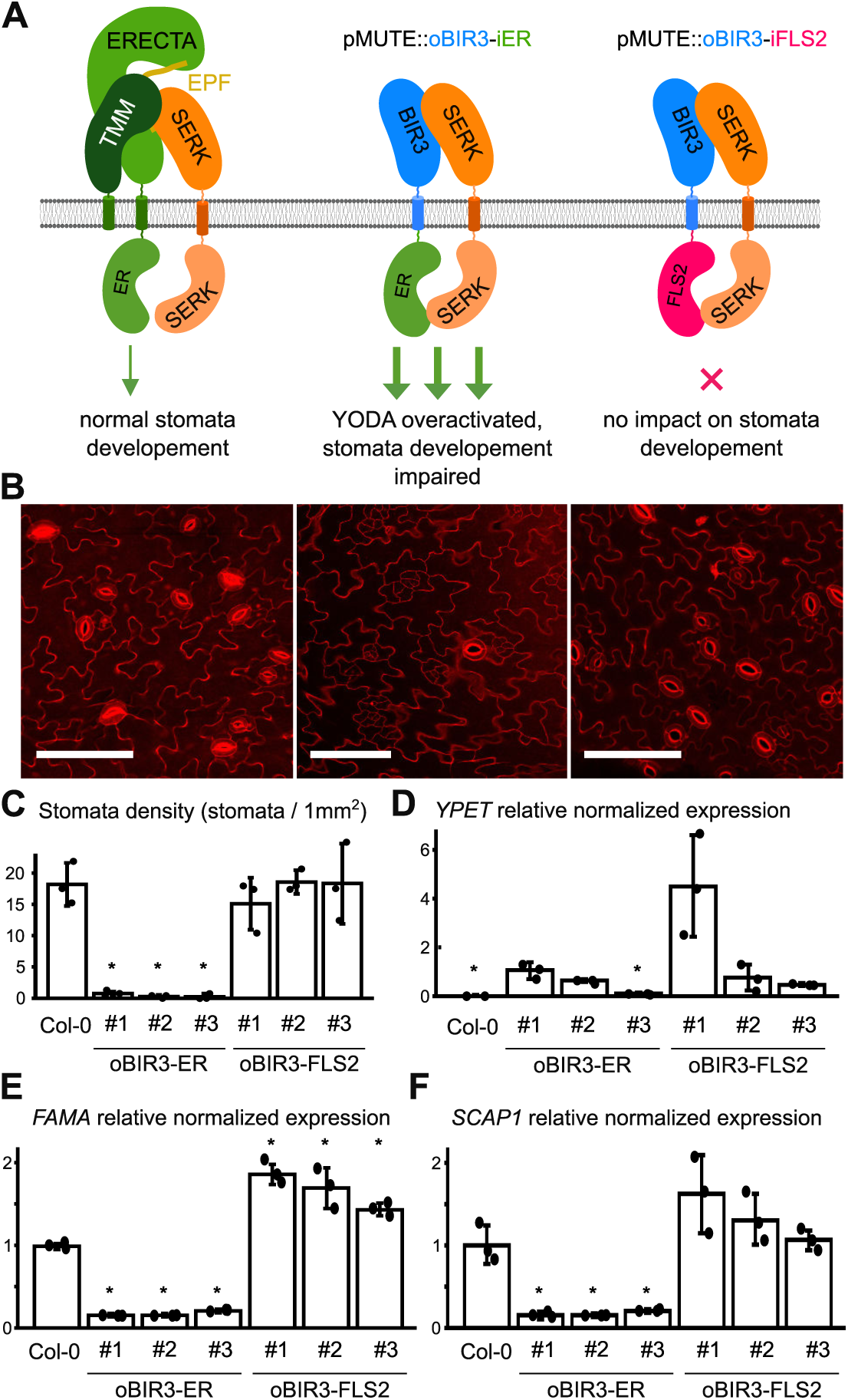
BIR3 – chimera reveal a conserved receptor activation mechanism in the LRR-RK ERECTA. **(A)** Schematic overview of ectopically expressed BIR chimera. The receptor kinase ERECTA interacts with SERK-co-receptor kinases upon ligand (EPF) binding and regulates stomata development (left). Expression of a oBIR3-iER chimera in the epidermis under the pMUTE promoter leads to pathway overactivation and the loss of stomata (middle), while the expression of an oBIR3-iFLS2 chimera has no effect on stomata development. **(B)** Confocal microscopy images of propidium iodide stained epidermis of the indicated genotype. Representative images of Col-0 (left panel), BIR3-ER-YPET (center), and BIR3-FLS2-YPET (right) are shown. Scale bar = 100 µm. **(C)** Abaxial stomata density of cotyledons (# numbers indicate independent lines). The average value of stomata density for three individual plants of each transgenic line is shown. Error bars depict standard deviations. Individual data points are shown as dot. Significant differences to wild type are indicated by an asterisk (t-test; p<0.05). **(D)** Expression level of the respective transgenes detected by qRT-PCR on YPET. The average values of three replicates are shown with error bars indicating standard deviations. Individual data points are shown as dots. Expression in the oBIR3-iER-YPET line #1 was arbitrarily set to 1. Significant differences in transgene expression to line #1 is indicated by an asterisk (t-test; p<0.05). **(E)** Relative normalized expression of *FAMA.* Normalized expression values of *FAMA* determined by quantitative RT-PCR are shown as average of three replicates. Error bars depict standard deviations. Individual data points are shown as dots. Expression in wild type was arbitrarily set to 1. Significant differences to wild-type levels are indicated by an asterisk (t-test; p<0.05). **(F)** Relative normalized expression of *SCAP1.* Normalized expression values determined by quantitative RT-PCR are shown as average of three replicates. Error bars display standard deviations. Individual data points are shown as dots. Expression in wildtype was arbitrarily set to 1. Significant differences to wild-type levels are indicated by an asterisk (t-test; p<0.05).

For each construct, we selected three representative lines according to YPET expression and measured density of mature stomata on cotyledons. The oBIR3-iER lines showed a drastic reduction of mature stomata and an increase in meristemoid-like cells on the leave surface (Figure 4B, C). In contrast, none of the oBIR3-iFLS2 lines showed any significant deviation from the wild-type phenotype, despite being expressed at a similar or higher level than the *BIR3-ER* chimeras (Figure 4D).

To analyze the observed phenotype on a molecular level, we tested expression of the guard mother cell (GMC)-specific transcription factor *FAMA* (Ohashi-Ito and Bergmann, 2006) and guard cell-specific Dof-type transcription factor *SCAP1* (Negi et al., 2013). The three independent BIR3-ER lines displayed a strong reduction of *FAMA* and *SCAP1* expression (Figure 4E,F), suggesting that the abnormal epidermal cells could be arrested at the meristemoid stage and do not express GMC-specific or guard cell-specific genes. All three oBIR3-iFLS2 expressing lines did not show a reduction in FAMA or SCAP1 expression. While SCAP1 transcript levels did not differ significantly from wildtype, there was a significant upregulation of FAMA expression in these lines (Figure 4E, F).

Finally, we tested if fusion of the BIR3 ectdomain to the LRR-RK GSO1/SGN3 (Tsuwamoto et al., 2008; Pfister et al., 2014) could restore the apoplastic barrier defects in the *sgn3-3* mutant (Pfister et al., 2014). GSO1/SGN3 directly senses the peptide ligands CASPARIAN STRIP INTEGRETY FACTORS 1 and 2 (CIF1, CIF2) to ensure proper formation of the Casparian strip, an endodermal diffusion barrier enabling selective nutrient uptake in the root (Pfister et al., 2014; Nakayama et al., 2017; Doblas et al., 2017; Okuda et al., 2020). A biochemical interaction screen has recently identified SERK proteins as putative co-receptor kinases for GSO1/SGN3 (Okuda et al., 2020), but it is presently unclear if SERKs mediate GSO1/SGN3 receptor activation *in vivo* (Figure 5A). We generated oBIR3-iSGN3, oBIR3-iSGN3^F146A,R170A^ and oBIR3-iSGN3^D1102N^ protein chimera expressed under the control of the *pSGN3* promoter in the *sgn3-3* mutant background. As previously described, the *sgn3-3* mutant has a non-functional apoplastic barrier that can be visualized and quantified by visualizing the uptake of the apoplastic tracer propidium iodide (PI) along the root and its access to the central vasculature (Figure 5B,C). We found that oBIR3-iSGN3 but none of the control lines restored *sgn3-3* apoplastic defects (Figure 5C, D), indicating a SERK-mediated GSO1/SGN3 receptor activation mechanism in the Casparian strip formation.

**Figure 5.**
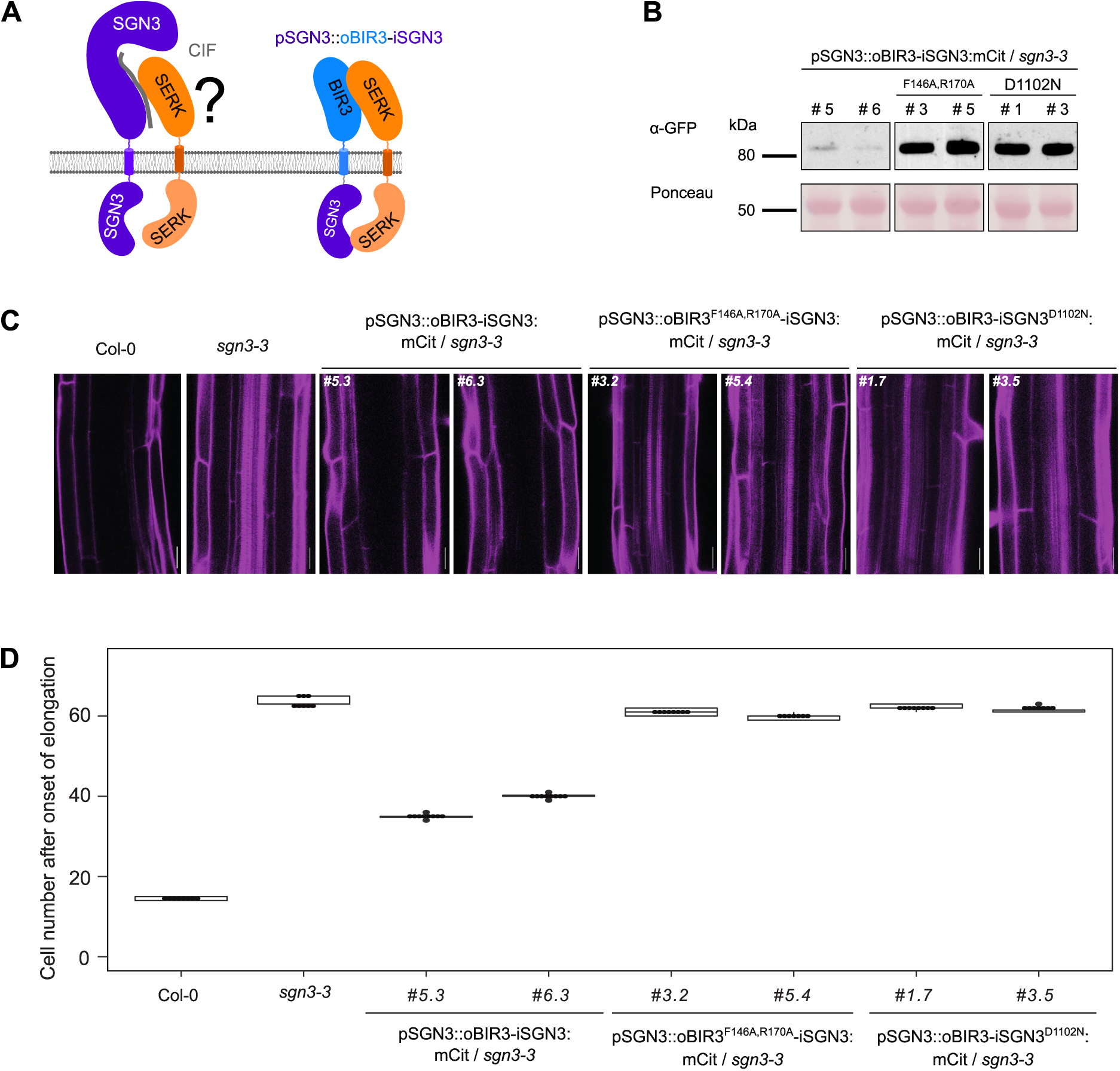
oBIR3-iSGN3 chimera suggest a role for SERK proteins in Casparian strip formation. **(A)** Schematic overview of a biochemically defined SGN3 – CIF – SERK signaling complex. The oBIR3-iSGN3 chimera is shown alongside. **(B)** Anti-GFP western blot on oBIR3-iSGN3 chimera in *sng3-3* background, with the Ponceau – stained membranes shown alongside. **(C)** Complementation of *sgn3-3* endodermal barrier defect by the chimera *SGN3::oBIR3-iSGN3.* Visualization of endodermal defects with the apoplastic tracer PI reaching the stele in barrier defective plants and blocked at the endodermis in plants with functional barriers. Pictures were taken around the 50th endodermal cell from the onset of elongation. Scale bar, 20 µm. **(D)** Quantification of PI block, measured as the number of endodermal cells after the onset of elongation where the PI block is observed. Data are presented as box plots with dot plot overlaid (n≥7). Different letters indicate significant differences between genotypes (p < 0.05).

## Discussion

The identification of a constitutive, ligand-independent interaction between the LRR ectodomains of two plant membrane signaling proteins prompted us to investigate if protein chimera between the BIR3 ectodomain and the cytoplasmic domain of various receptor kinases would lead to constitutively active signaling complexes. Despite the significant structural differences between LRR-RK – SERK and BIR – SERK complexes, our data demonstrates that a wide range of oBIR3 – iLRR-RK chimera are functional *in planta*.

Expression of the oBIR3 – iBRI1 chimera resulted in a strong, constitutive activation of the brassinosteroid signaling pathway. The gain-of-function effect is much stronger than previously described for the BRI1 *sud1* and SERK3 *elongated* alleles, respectively (Belkhadir et al., 2012; Jaillais et al., 2011; Hohmann et al., 2018a), and comparable to constitutive activation of BES1 (Figure 2A) (Yin et al., 2002). The constitutive signaling activity of the oBIR3 – iBRI1 chimera depends on the ability of the BIR3 ectodomain to bind SERK ectodomains and on the kinase activity of the BRI1 cytosolic segment (Figure 2). This reinforces the notion that formation of the heterodimeric extracellular signaling complex drives LRR-RK receptor activation, and that signaling specificity is encoded in the kinase domain of the receptor, not the co-receptor (Bojar et al., 2014; Hohmann et al., 2018b; Zheng et al., 2019). The phenotypes of oBIR3 – iBRI1 and *bes1-1D* plants in addition suggest that little signal amplification appears to occur throughout the brassinosteroid signaling pathway (Figure 2).

Analysis of the oBIR3 – iHAE chimera revealed a strongly conserved activation mechanism between different, SERK-dependent LRR-RK signaling pathways, as previously suggested (Hohmann et al., 2018b) (Figure 3). In addition, our experiments suggest that BIR ectodomains are able to interact with SERK proteins in the abscission zone, and thus BIR proteins may act as negative regulators of HAESA / HSL2 mediated signaling cascades in wild-type plants (Figure 3). In this respect, it is noteworthy that the BIR suppressor SoBIR1/EVERSHED has been previously characterized as a genetic component of the floral abscission signaling pathway (Leslie et al., 2010).

ERECTA family kinases have been previously shown to require SERK co-receptor kinases to control stomatal patterning and immune responses (Meng et al., 2015; Jordá et al., 2016). Our functional oBIR3 – iER chimera now suggests, that, despite the requirement for TMM, EPF bound ER signaling complexes are activated by SERK proteins in very similar ways as previously reported other LRR-RKs (Hohmann et al., 2017) (Figure 4). Expression of the oBIR3 – iER chimera in meristemoid cells lead to a similar phenotype as described for the expression of constitutively active versions of YODA, MKK4, and MKK5 (Lampard et al., 2009). This strongly indicates that the oBIR3 – iER chimera displays constitutive, ligand-independent signaling activity. The specificity of signal transduction seems to be largely maintained, as expression of oBIR3 -iFLS2 led to wild-type like stomatal development. At the molecular level, however, we observed a significant increase of *FAMA* expression in all tested oBIR3 – iFLS2 lines. This is consistent with an antagonistic regulation of these two pathways (Sun et al., 2018). The observed upregulation of *FAMA* expression, however, did not significantly alter stomata density. This is likely because the transcriptional activation of the oBIR3-iFLS2 construct in this experiment happens only in meristemoid cells and might be compensated by post-transcriptional regulation.

Finally, our finding that expression of a oBIR3 – iSGN3 chimera could partially rescue the apoplastic barrier defects of the *sgn3-3* mutant. Notably, BIR ectodomains specifically bind the ectodomains of SERKs (Ma et al., 2017; Hohmann et al., 2018a), while not forming complexes with the LRR ectodomain of the sequence-related NSP-INTERACTING KINASE1 (NIK1) (Figure 6). This suggests that SERK proteins may have redundant functions in SGN3/GSO1 signaling in the endodermis (Pfister et al., 2014; Okuda et al., 2020).

**Figure 6.**
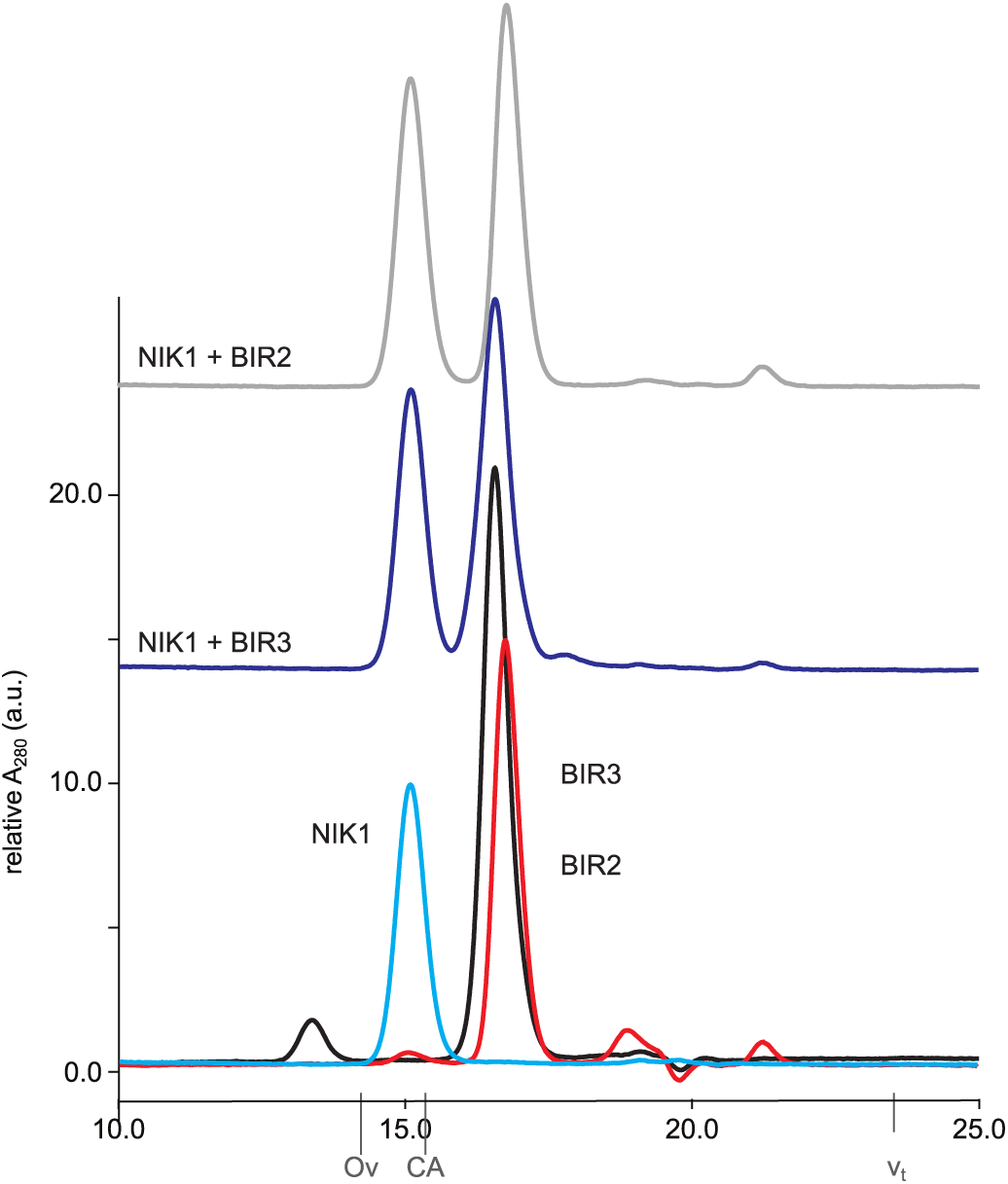
The LRR-ectodomains of BIRs and NIK1 do not interact *in vitro*. Analytical size-exclusion chromatography binding experiments using the NIK1, BIR2 and BIR3 ectodomains. BIR2 (gray absorption trace) and BIR3 (in dark blue) form no complex with NIK1, their respective elution volumes correspond to that of the isolated protein (BIR2 in red, BIR2 in black, NIK1 in light blue). The NIK1 LRR domain shares 49% amino-acid sequence identify with the SERK1 ectodomain. The total volume (*v*_t_) is shown together with elution volumes for molecular mass standards (Ov, Ovalbumin, 44,000 Da; CA, Carbonic anhydrase, 29,000 Da).

Taken together, the simple, lego-style assembly of BIR3 chimera (Figure 7) and the availability of suitable control lines now enables the genetic characterization of orphan LRR-RKs with unknown/unclear loss-of-function phenotypes and dissection of their potential activation mechanism. BIR3 protein chimera may also be of use for biochemical or genetic interaction screens, in which a constitutively active form of the receptor is desirable.

**Figure 7.**
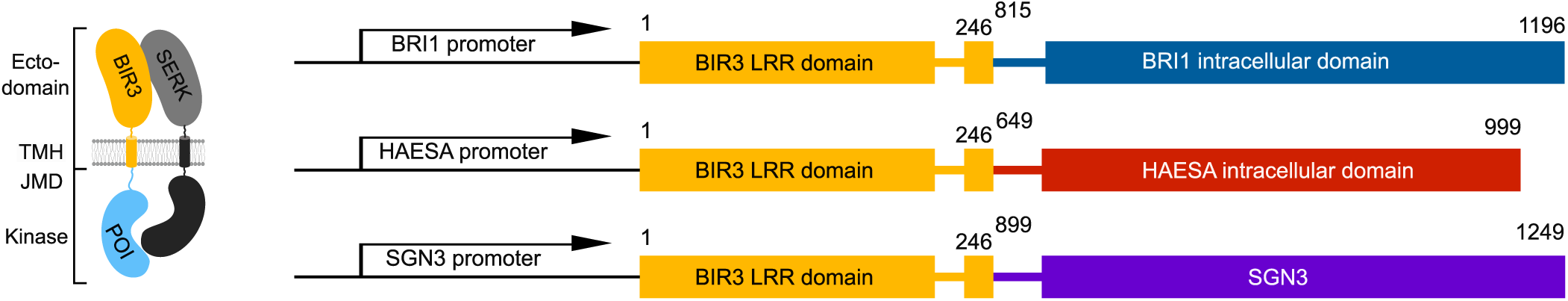
Design principles of BIR chimeras. Schematic overview of selected BIR3 chimera used in this study. Chimeric proteins are expressed under the endogenous promoter of the respective receptor.

## Material & Methods

### Plant material, growth conditions and generation of transgenic lines

To design chimeric receptor kinases, the transmembrane helix of all LRR-RKs was first predicted using TMHMM (version 2.0, https://services.healthtech.dtu.dk/service.php?TMHMM-2.0) (Krogh et al., 2001). The native signal peptide, extracellular domain and the transmembrane helix from AtBIR3 (residues 1-246) were fused to the juxtamembrane and kinase domains of the respective receptor (BRI1 residues 815-1196, HAE 649-999, SGN3 899-1249). No additional linker sequences were added (Figure 7). Fragments were amplified from *Arabidopsis thaliana* (ecotype Col-0) genomic or cDNA and cloned into pDONR221 (ThermoFisher Scientific) using Gibson-cloning technology; mutations were introduced through site directed mutagenesis (Supplemental Table 1). Binary vectors were assembled using the multi-site Gateway technology into the binary vector pB7m34GW, harboring a Basta resistance gene (ThermoFisher Scientific). All constructs were introduced into *Agrobacterium tumefaciens* strain pGV2260, and *Arabidopsis* plants were transformed using the floral dip method (Clough and Bent, 1998).

GABI_134E10 was used as a *bri1*-null allele (Jaillais et al., 2011), ABRC CS65988 as *bes1-1D* (Yin et al., 2002), ABRC CS6159 as *det2-1* (Chory et al., 1991), and *hae hsl2* and *sgn3-3* as previously reported (Stenvik et al., 2008; Pfister et al., 2014). All plants were grown in 50 % humidity, 21 °C and a 16 h light – 8 h dark cycle.

To generate the chimeric *pMUTE::BIR3-FLS2-Ypet* and *pMUTE::BIR3-ERECTA-Ypet* genes, a 1946 bp DNA fragment comprising the coding sequence of the N-terminal extra-cellular domain of BIR3 (residues 1-245) followed by a short multiple cloning site, the coding sequence of YPET, and a 411 bp terminator sequence of the Arabidopsis *UBQ10* gene was synthesized (Baseclear, The Netherlands) and inserted in the T-DNA of a modified pCambia3300 binary vector. A 2432 bp promoter region of the Arabidopsis *MUTE* gene was PCR amplified from Col-0 genomic DNA and inserted directly upstream of the synthetic BIR3 fusion construct by in-fusion cloning (Clonetech). The coding regions for the intracellular domains of FLS2 (residues 807-1173) and ERECTA (residues 581-976) were PCR amplified from Arabidopsis seedling-derived cDNA and inserted in-frame between the coding region of the BIR3 extracellular domain and the YPET coding region. All constructs were confirmed by Sanger sequencing.

### Hypocotyl growth assay

Seeds were surface sterilized, stratified at 4 °C for 2 d, and plated on ½ MS, 0.8 % agar plates supplemented with 1 μM brassinazole (BRZ, from a 10 mM stock solution in 100 % DMSO, Tokyo Chemical Industry Co. LTD) or, for the controls, with 0.1 % (v/v) DMSO. Following a 1 h light exposure to induce germination, the plates were wrapped in aluminium foil and incubated in the dark at 22 °C for 5 d. The plates were then scanned at 600 dpi on a regular flatbed scanner (CanoScan 9000F, Canon), hypocotyl lengths measured using Fiji (Schindelin et al., 2012) and analyzed in R (R Core Team, 2014) (version 3.6.1) using the packages mratios (Kitsche and Hothorn, 2014) and multcomp (Hothorn et al., 2008). Rather than p-values, we report unadjusted 95% confidence limits for fold-changes. A mixed effects model for the ratio of of a given line to the wild-type Col-0, allowing for heterogeneous variances, was used to analyze log-transformed endpoint hypocotyl lengths. To evaluate the treatment-by-mutant interaction, the 95 % two-sided confidence intervals for the relative inhibition (Col-0: untreated vs. BRZ-treated hypocotyl length)/(any genotype: untreated vs. BRZ-treated hypocotyl length) was calculated for the log-transformed length.

### Plant protein extraction and immunoprecipitation

Seeds were plated on ½ MS, 0.8 % agar plates and grown for ∼ 14 d after surface sterilization and stratification. Seedlings were harvested, padded dry carefully, snap-frozen in liquid N_2_, and ground to fine powder using pre-cooled mortar and pestel. 1 g of powder per sample was resuspended in 3 ml of ice cold extraction buffer (50 mM Bis Tris pH 7.0, 150mM NaCl, 10 % (v/v) glycerol, 1 % Triton X-100, 5 mM DTT, protease inhibitor cocktail (P9599, Sigma)) and agitated gently at 4 °C for 1 h. Subsequently, samples were centrifuged (30 min, 16,000 g, 4 °C), the supernatant then transferred to a fresh tube and the protein concentration estimated through a Bradford assay with a BSA standard curve.

For each co-immunoprecipitation (Co-IP), 20 mg of total protein in a volume of 5 ml were incubated with 50 μl of anti-GFP superparamagnetic MicroBeads (Miltenyi Biotec) for 1 h at 4 °C with gentle agitation. Using a magnetic rack and μMACS Columns (Miltenyi Biotec) which were washed once with extraction buffer the beads were collected and then washed 4 times with 1 ml of ice cold extraction buffer. Bound proteins were then eluted in 2 times 20 μl of extraction buffer pre-heated to 95 °C. Samples were then separated on 10 % SDS-PAGE gels and analyzed with a standard western blot using the following antibodies: anti-GFP antibody coupled to horse radish peroxidase (Anti-GFP-HRP, Miltenyi Biotec 130-091-833) at 1:2,000 dilution to detect mCitrine; anti-SERK3 (Bojar et al., 2014) at 1:5,000 dilution in conjunction with a secondary anti-rabbit HRP antibody (1:10,000, Calbiochem #401353) to detect SERK3.

### Western blot for BES1

For each sample, ∼ 100 µg of seven day old seedlings, grown on ½ MS, 0.8 % agar plates, were harvested, frozen in liquid N2 and ground to powder using bead mill (Retsch MM400). The sample was resuspended in ∼ 200 µl of ice cold extraction buffer (25 mM Tris pH 7.5, 150 mM NaCl, 1 % SDS, 10 mM DTT, protease inhibitor cocktail (P9599, Sigma)), incubated with gentle agitation for 1 h at 4 °C, centrifuged for 30 min at 4 °C, 16,000 g. The supernatant was transferred to a fresh tube and the protein concentration assessed through a Bradford assay. 80 µg of total extracted protein were separated on a 12 % SDS-PAGE gel and analyzed in a westernblot (primary antibody: anti-bes1, 1:2,000 (Yin et al., 2002), secondary antibody: anti-rabbit HRP (1:10,000, Calbiochem #401353)).

### Stomata density measurements and microscopy

Seven-day old T2 seedlings were used to determine stomata density. For confocal imaging, seedlings were incubated in 10mg/L propidium iodide (PI) solution for 30 min, and then washed with water. Abaxial epidermal regions of cotyledons were imaged using a Zeiss LSM 780 NLO microscope equipped with a Plan-Apochromat 25x/0.8 Imm Korr DIC objective. PI staining was visualized with an excitation wave length of 514 nm and emission was recorded between 566 nm and 643 nm. Mature stomata were counted in a 0.25 mm by 0.25 mm epidermal area for three seedlings of each line.

### Expression analysis

Seven-day old T2 seedlings were used to analyze expression levels of the transgene as well as endogenous genes. For each independent line, RNA was extracted from 24 pooled T2 seedlings using the RNase® Plant Mini Kit (QiaGen). cDNA was synthesized using the RecertAid First Strand cDNA Synthesis Kit (Thermo Scientific). Relative abundance of the endogenous *FAMA* and *SCAP1* transcripts as well as chimeric *YPET*-containing *BIR3* transcripts were measured by quantitative RT-PCR (program: 1. 50°C for 10 min, 2. 95°C for 5 min, 3. 95°C for 10 s, 4. 60 °C for 30 s; repeat step 3 – 4 40 times; 5. 95°C for 10 s, 6. ramp 65°C to 95 and increase 0.5°C every 5s, Plate Read). Expression levels of endogenous *ACTIN2* were used for normalization.

### Propidium iodide permeability assay and confocal microscopy of wild-type and complemented *sgn3-3* plants

Propidium Iodide (PI) permeability assay were performed on 5 d old seedlings. In brief, the seedlings were stained in dark for 10 mins in 10µg/ml PI, rinsed twice in water and quantified as previously described (Naseer et al., 2012). Endodermal cell numbers were quantified using a Leica Epifluorescence microscope. Representative confocal images were acquired with a Leica SP8, with excitation and detection windows set as follows for PI: excitation −488 nm, emission – 500-550 nm. Confocal images were processed and analyzed using ImageJ. For samples treated with the CIF2 peptide, the seedlings were grown on ½ MS for 3 days followed by transfer to ½ MS + 10µM CIF2 peptide for 2 d and subsequently analyzed for PI permeability. Statistical analyses were done in the R environment (R Core Team, 2014). For multiple comparisons between genotypes, Kruskal-Wallis’ test was performed and nonparametric Tukey’s test was subsequently used as a multiple comparison procedure. Different letters indicates significant difference (P<0.05). Data are presented as box plots overlaid with dot plots.

### Protein expression, purification and size exclusion chromatography

The coding sequence of AtNIK1^32-248^ was amplified from *Arabidopsis thaliana* cDNA, AtBIR2^1–222^, BIR3^1–213^ from *A. thaliana* genomic DNA and fragments were cloned into a modified pFastBac vector (Geneva Biotech), providing a tobacco etch virus protease (TEV)-cleavable C-terminal StrepII-9xHis tag. NIK1 was fused to an N-terminal azurocidin secretion peptide. Proteins were expressed by infection of *Trichoplusia ni* (strain Tnao38) (Hashimoto et al., 2010) cells with 15 ml of virus in 250 ml of cells at a density of ∼ 2× 10^6^ cells ml^-1^, incubated for 26 h at 28 °C and 110 rev min^-1^ and then for another 48 h at 22 °C and 110 rev min^-1^. Secreted proteins were purified from the supernatant by sequential Ni^2+^ (HisTrap excel; GE Healthcare; equilibrated in 25 mM KP_i_ pH 7.8, 500 mM NaCl) and StrepII (Strep-Tactin XT; IBA; equilibrated in 25 mM Tris pH 8.0, 250 mM NaCl, 1 mM EDTA) affinity chromatography followed by size-exclusion chromatography on a HiLoad 16/600 Superdex 200pg column (GE Healthcare), equilibrated in 20 mM sodium citrate pH 5.0, 250 mM NaCl. The theoretical molecular weight of the purified ectodomains is 23.6 kDa for AtNIK1^32-248^, 23.4 kDa for AtBIR2^1–222^ and 24.0 kDa for BIR3^1–213^.

For analytical size exclusion chromatography experiments, a Superdex 200 increase 10/300 GL column (GE Healthcare) was pre-equilibrated in 20 mM sodium citrate pH 5.0, 250 mM NaCl. For each run, 40 μg of the individual NIK1, BIR2 or BIR3 ectodomains were injected in a volume of 100 μl and elution at 0.75 ml min^-1^ was monitored by ultraviolet absorbance at λ = 280 nm. To probe interactions between NIK1, BIR2 and BIR3, 40 μg of the respective proteins were mixed in a total volume of 100 μl and incubated on ice for 30 min before analysis as outlined above.

## SUPPLEMENTAL DATA

### Supplemental figures

**Supplemental Figure 1.**
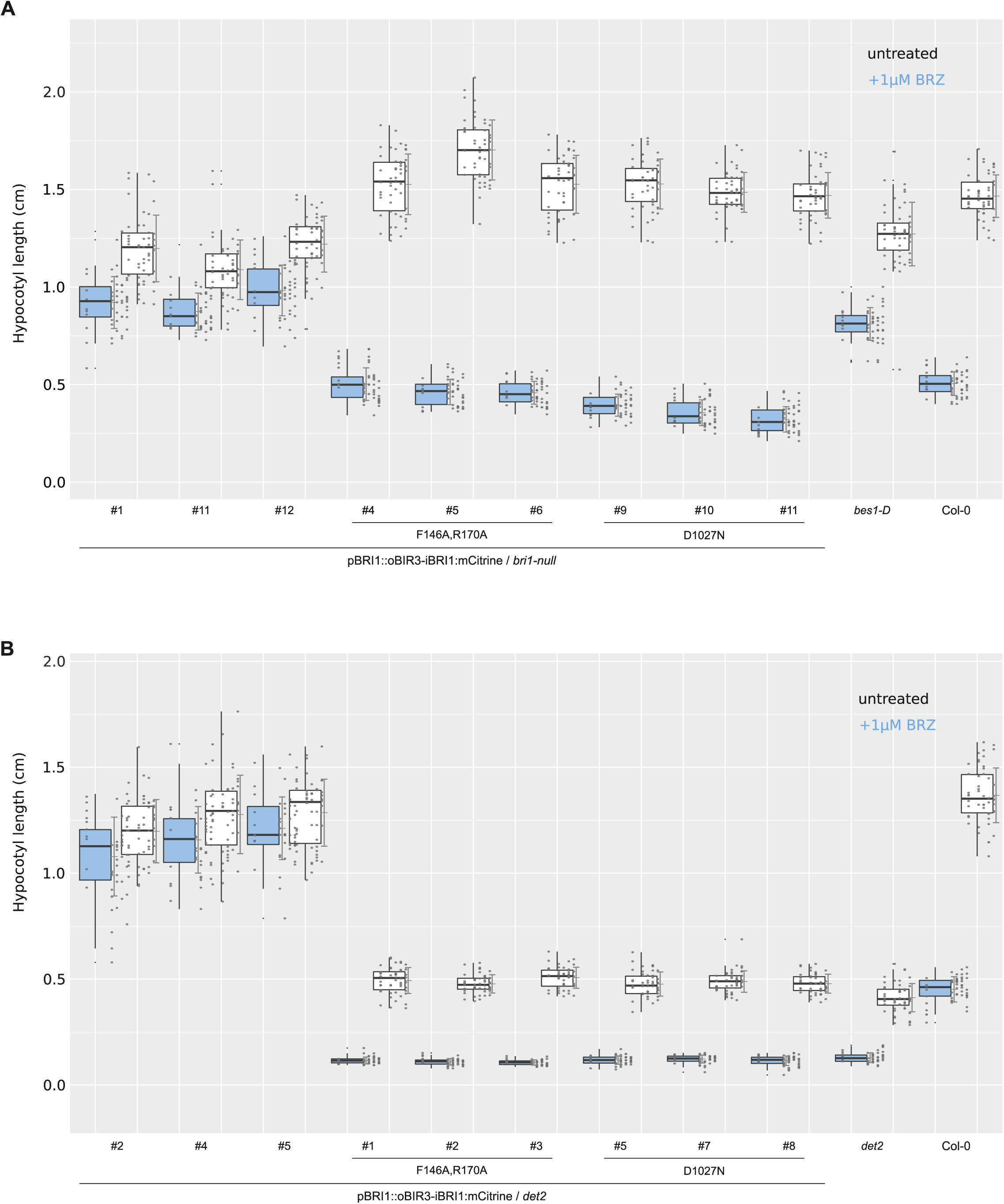
Hypocotyl growth assay raw data (supports Figure 2A). Depicted are box plots (center line, median; box limits, upper and lower quartiles; whiskers, 1.5x interquartile range; points, outliers) together with the raw data (depicted as individual dots, grouped per plate) and mean ± standard deviation alongside. The raw data for oBIR3-iBRI1 chimera in the *bri1-null* background is shown in (A) and in the *det2* – background in (B). Untreated: white, BRZ treated: blue. For each sample n= 50 biologically independent hypocotyls, coming from 5 different ½MS plates, have been measured.

**Supplemental Figure 2.**
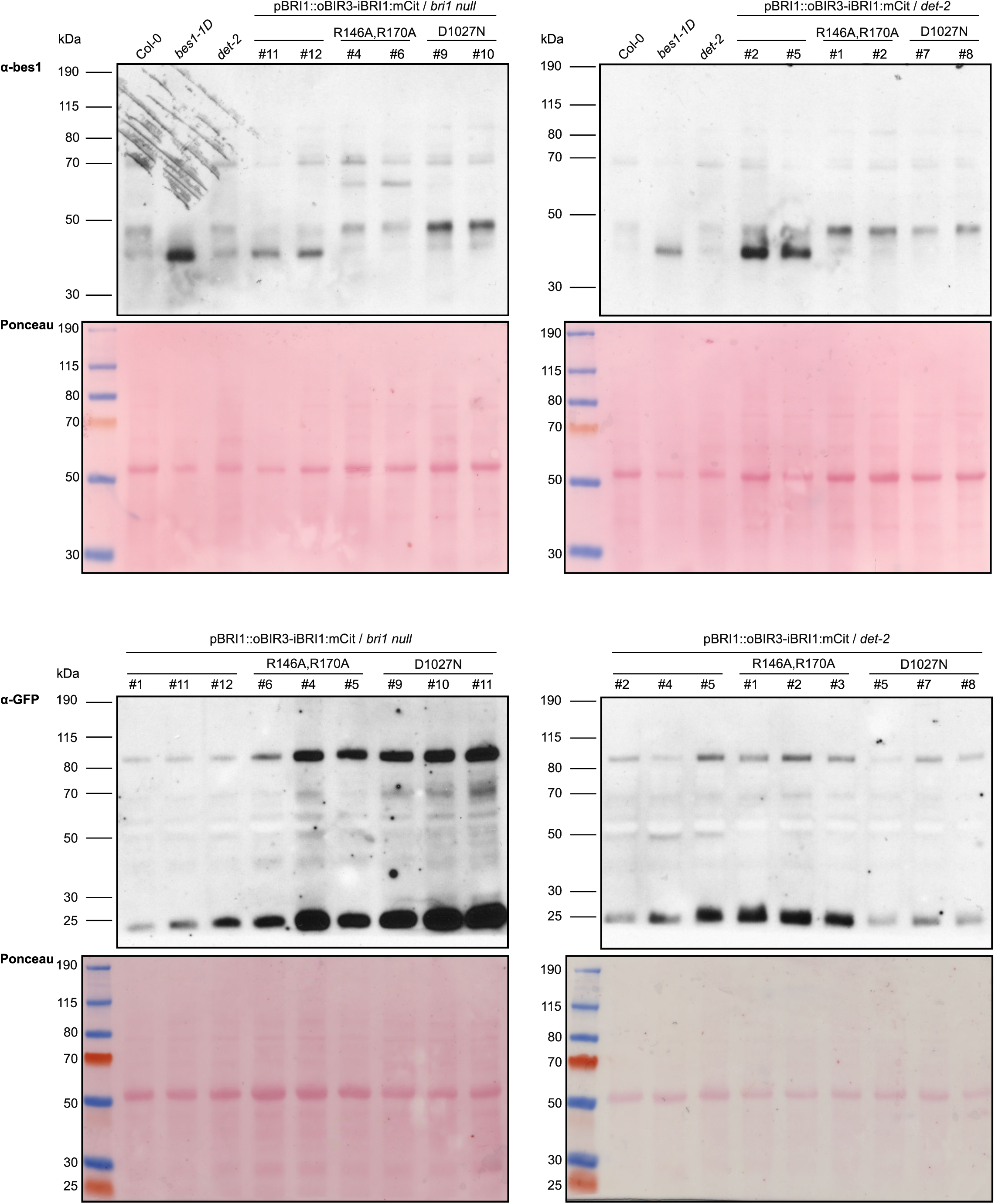
Full western blot films and Ponceau – stained membranes (supports Figure 2A-C). Scans of the full western blot films and the Ponceau - stained membranes used to prepare Figures 2 A – C are shown.

**Supplemental Figure 3.**
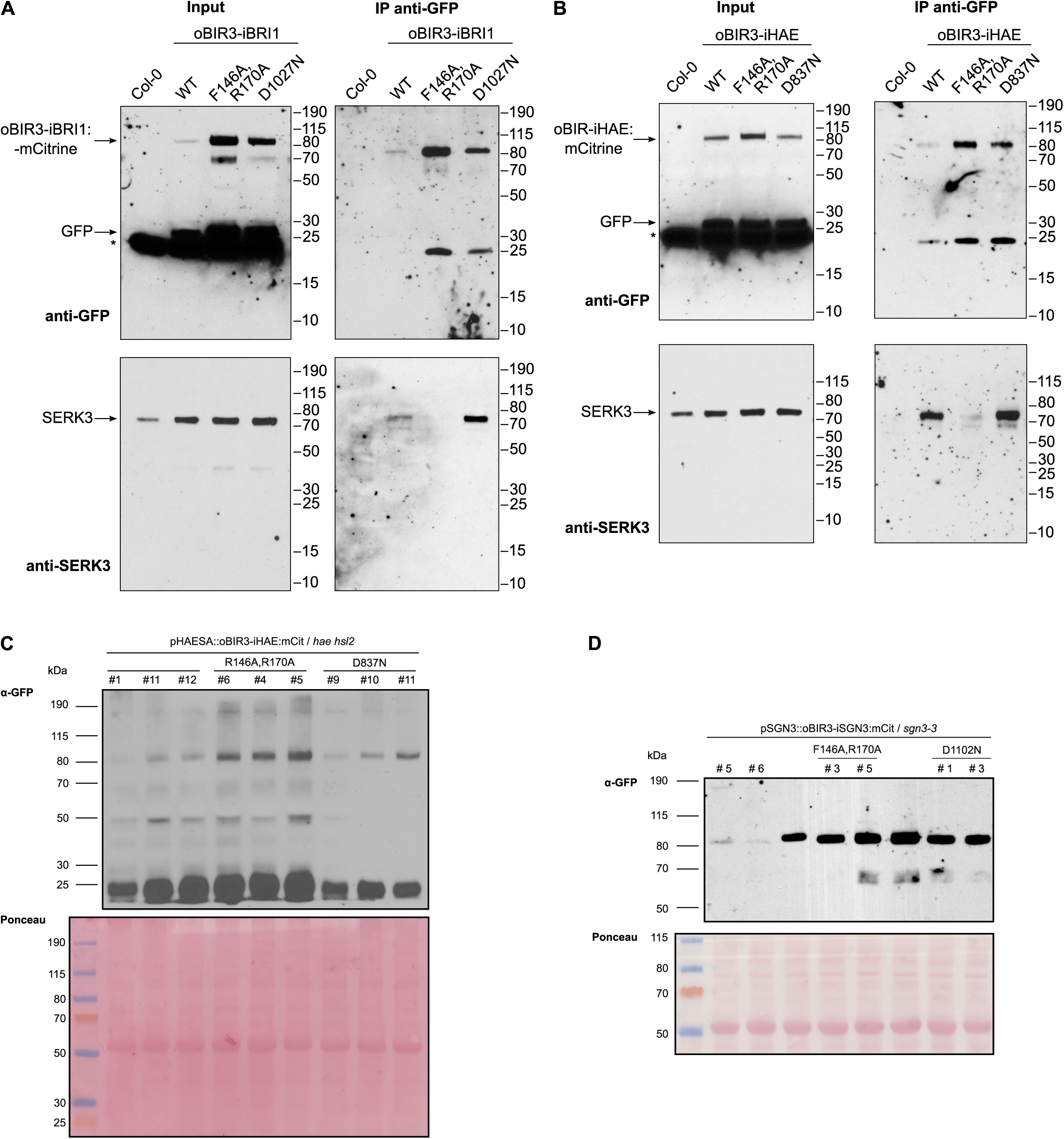
Full western blot films and Ponceau – stained membranes (supports Figure 2D and 3C-D). Scans of the full western blot films and the Ponceau - stained membranes used to prepare Figures 2D as well as Figure 3 C-D are shown.

**Supplemental Table 1:**
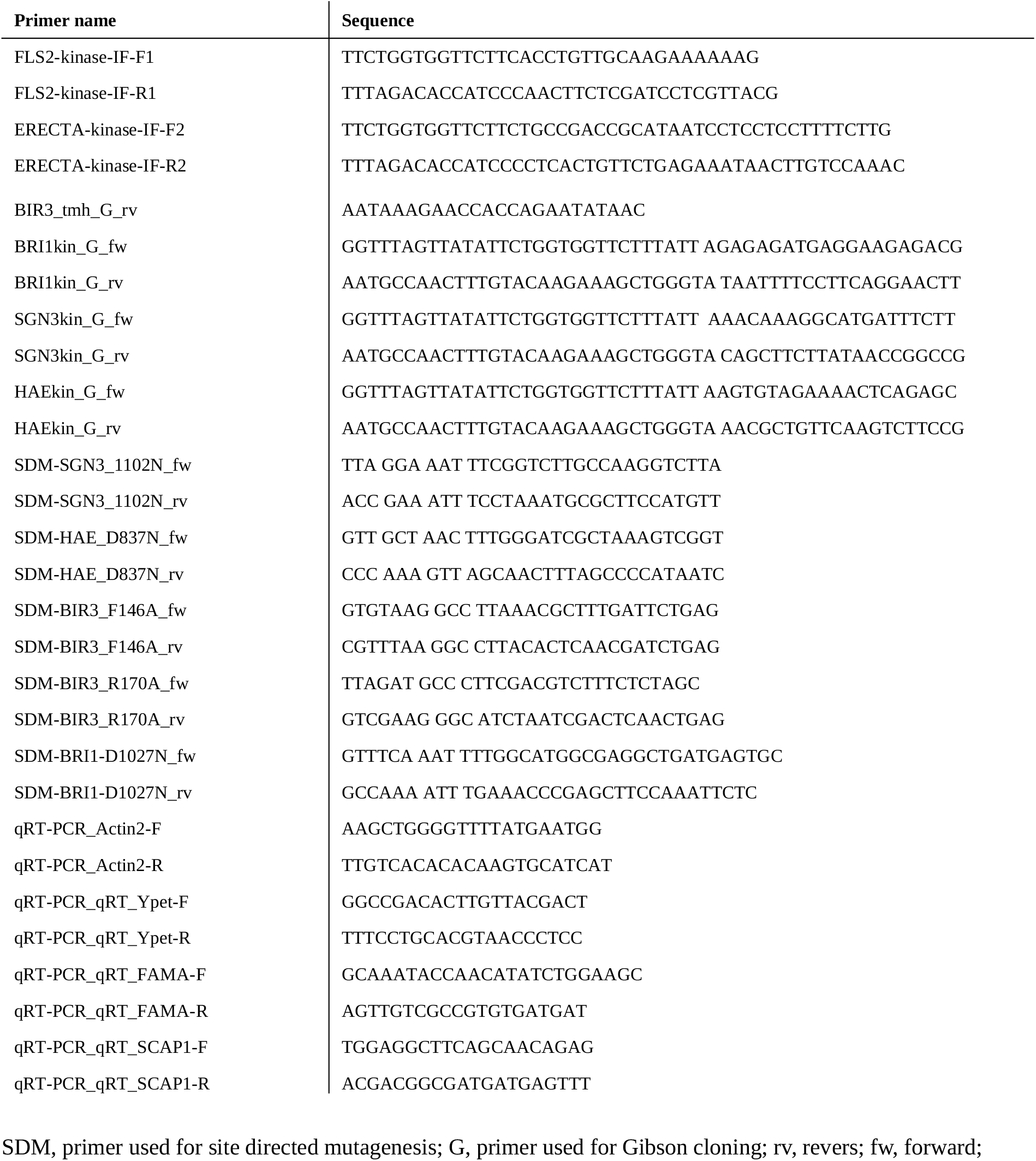
Primers used in this study.

